# Freeze-tolerant crickets fortify their actin cytoskeleton in fat body tissue

**DOI:** 10.1101/2024.11.22.624896

**Authors:** Maranda L. van Oirschot, Jantina Toxopeus

## Abstract

Animals that overwinter in temperate climates must prevent or repair damage to their cells to survive winter, but we know little about how they protect cellular structure at the cytoskeletal level. Both chilling (no ice formation) and freezing (ice formation) are hypothesized to cause substantial challenges to cell structure and the cytoskeleton. The spring field cricket *Gryllus veletis* becomes freeze-tolerant following a 6-week acclimation to fall-like conditions, during which they differentially express multiple cytoskeleton-related genes. We tested the hypothesis that *G. veletis* alter their cytoskeleton during acclimation to support maintenance of cytoskeletal structure during freezing and thawing. We used immunocytochemistry and confocal microscopy to characterize changes in microfilaments (F-actin, a polymer of G-actin) and microtubules (a polymer of α- and β-tubulin) in three tissues. While we saw no effect of acclimation on microtubules, crickets increased the abundance of microfilaments in fat body and Malpighian tubules. When we chilled or froze these freeze-tolerant crickets, there was no apparent damage to the actin or tubulin cytoskeleton in fat body, but there was decreased cytoskeleton abundance in Malpighian tubules. When we froze freeze-intolerant (unacclimated) crickets, microfilament abundance decreased in fat body tissue, while microfilaments were unaffected by chilling to the same subzero temperature. Our study shows that freeze-tolerant crickets are able to prevent or rapidly repair ice-induced damage to the actin cytoskeleton in fat body, likely due to preparatory changes in advance of freezing – i.e., during acclimation. We suggest future directions examining the mechanisms that underlie these structural changes.

**Summary Statement:** Freeze-tolerant crickets (*Gryllus veletis*) modify their actin cytoskeleton in select tissues to prevent freeze-induced cellular damage, while freeze-intolerant crickets of the same species experience irreparable cytoskeleton damage during freezing.

## Introduction

Low temperatures pose many challenges for organisms, and it is increasingly important to understand the cellular and physiological mechanisms that facilitate survival in cold environments as extreme low temperature events become more frequent due to climate change (Marshall et al., 2020; Williams et al., 2015). Most insects are chill-susceptible or cold-intolerant, and are killed by mild low temperatures (e.g., above 0°C), while cold-tolerant insects can survive subzero temperatures by being freeze-avoidant or freeze-tolerant (Overgaard and MacMillan, 2017; Toxopeus and Sinclair, 2018). Freeze-avoidant organisms physiologically depress the temperature at which ice formation begins - known as the supercooling point (SCP) - but die if ice forms (Lee, 2010; Toxopeus and Sinclair, 2018). Conversely, freeze-tolerant organisms survive at least 50% of their body liquid turning to ice (Storey and Storey, 2013; Toxopeus and Sinclair, 2018). Ice can cause substantial cellular injury via mechanical damage to membranes, freeze-induced cellular dehydration, and impacts on metabolism (Lee, 2010; Toxopeus and Sinclair, 2018). Freeze-tolerant insects must either be able to prevent this cellular injury, or successfully repair nonlethal injury post-freezing (Teets et al., 2023; Toxopeus and Sinclair, 2018). Many of the proposed mechanisms underlying freeze tolerance are correlated with seasonal changes in cold tolerance. For example, acclimatization to fall conditions (or acclimation to similar settings in a laboratory) can stimulate accumulation of cryoprotectant molecules that protect cells at low temperatures (Koštál et al., 2011; Toxopeus et al., 2019a). While cryoprotectants have been well-studied, the cytoskeleton is gaining increased attention as a potentially important component of insect freeze tolerance (Des Marteaux et al., 2018a; Toxopeus and Sinclair, 2018; Toxopeus et al., 2019b), and will be the focus of our study in the freeze-tolerant cricket *Gryllus veletis* (Alexander & Bigelow) (Orthoptera: Gryllidae).

The cytoskeleton is a dynamic matrix of proteinaceous filaments that affects cell structure, cell motility, the movement of substances within cells, and cell-cell connections. Microfilaments form when globular (G)-actin proteins polymerize into filamentous (F)-actin (Cooper and Adams, 2000), while microtubules are polymers of α and β-tubulin subunit proteins (Cooper and Adams, 2000; Li and Moore, 2020). Polymerization and stability of microfilaments and microtubules requires ATP or equivalents (Cooper and Adams, 2000; Dominguez and Holmes, 2011). In addition, various cytoskeleton-binding and cytoskeleton-associating proteins are important for regulating the length and arrangement of these cytoskeletal fibers. For example, villins and gelsolins are involved in severing and capping actin microfilaments (Dominguez and Holmes, 2011; Pollard, 2016). Of particular relevance to this study, microfilament and microtubule structure relies on intermolecular interactions (i.e., between actin subunits, between α and β-tubulin subunits) that can be sensitive to cold (Li and Moore, 2020).

Low temperatures and freezing can disrupt microtubules and microfilaments, both during and after a cold or freezing stress. For example, microtubule depolymerization (dissociation into tubulin subunits) has been recorded during chilling (0-4°C) of tobacco cells (Pokorná et al., 2004) and yeast cells (Li and Moore, 2020) and during extracellular freezing (down to −6°C) of spinach cells (Bartolo and Carter, 1991). Microfilaments may also depolymerize during cold exposure (e.g., in tobacco cells at 0°C; Pokorná et al., 2004), and can be damaged by intracellular ice formation (e.g., in rapidly frozen human cells; Müllers et al., 2019), but seem to be more stable during cold and freezing stress than microtubules (Li and Moore, 2020; Müllers et al., 2019). Microtubule and microfilament depolymerization during cold and freezing does not appear to be driven by decreased abundance of the tubulin and G-actin subunits, respectively (Li and Moore, 2020; Pokorná et al., 2004), although cytoskeletal subunits may leak out of cells with substantial membrane damage (Ragoonanan et al., 2010). The destabilization of these cytoskeletal filaments is more likely driven by the impacts of cold and freezing on the intermolecular interactions between subunits, or reduced function of cytoskeletal cofactors and chaperones (Li and Moore, 2020). Indeed, both microfilaments and microtubules can successfully repolymerize when cold-shocked or frozen cells are returned to a warm temperature, often within seconds to minutes (Bartolo and Carter, 1991; Li and Moore, 2020; Pokorná et al., 2004). Cytoskeletal depolymerization and damage can also be observed after cells are rewarmed. For example, cold-intolerant spinach cells rapidly repolymerize microtubules post-thaw, but then exhibit further depolymerization within a day of ‘recovery’ (Bartolo and Carter, 1991). We hypothesize that cold-tolerant organisms protect their cytoskeleton (Toxopeus and Sinclair, 2018) by either 1) preventing depolymerization during cold and freezing, or 2) successfully repolymerizing and maintaining the repolymerized state of the cytoskeleton after cold or freezing.

Many organisms exhibit increased protection of the cytoskeleton in association with increased cold tolerance or freeze tolerance. For example, chill-susceptible fall field crickets (*Gryllus pennsylvanicus*) show an increase in cold hardiness and an increase in F-actin intensity in rectal epithelial cells following acclimation to mild low temperatures (12°C for 1 week; Des Marteaux et al., 2018b). The microfilaments of these cold-acclimated crickets are more resistant to depolymerization following cold shock (−4°C for 1 h) than unacclimated crickets (Des Marteaux et al., 2018b). Similarly, cold-acclimation (5°C for 1 month) of onion maggot (*Delia antiqua*) pupae protects microfilaments within Malpighian tubules following a more extreme cold shock of supercooling (not freezing) to −20°C for 12 days (Kayukawa and Ishikawa, 2009).

Acclimation can also help protect microtubules from cold and freezing, as seen in cells from spinach plants that were acclimated to 5°C for 2 weeks and could better maintain (resist depolymerization of) their microtubules after freezing and thawing than their unacclimated counterparts (Bartolo and Carter, 1991). Changes to the cytoskeleton may be promoted by several factors, including transcriptional upregulation of actin (Kim et al., 2006), tubulin (Des Marteaux et al., 2017), proteins that interact with the cytoskeleton (Des Marteaux et al., 2017; Kayukawa and Ishikawa, 2009; Ouellet et al., 2001; Zhang et al., 2011), and cold-specific isoforms of these proteins (Carrasco et al., 2011). However, some cold-tolerant insects downregulate actin (Des Marteaux et al., 2017) or tubulin (Kim and Denlinger, 2009), and may show minimal changes to their cytoskeleton in the cold- or freeze-tolerant state. For example, microtubules become less abundant in the thoracic (flight) muscle of *C. pipiens* mosquitoes during diapause, a state of enhanced cold tolerance (Kim and Denlinger, 2009). In addition, freeze-tolerant larvae of the drosophilid fly *Chymomyza costata* show very few differences in either microfilament or microtubule arrangement in multiple tissues compared to freeze-intolerant larvae, even after freezing and thawing (Des Marteaux et al., 2018a). Damage to the radial network of microtubules within fat body cells was detected in these flies after freezing, but repair of this damage post-thaw did not seem necessary for survival of the organism (Des Marteaux et al., 2018a). With the exception of *C. costata* (Des Marteaux et al., 2018a), we know little about the relationship between freezing and the cytoskeleton in naturally freeze-tolerant animals.

The distribution and abundance of cytoskeletal components differ among insect cell types. Many cells have a layer of cortical F-actin supporting the plasma membrane, e.g., fat body cells (Des Marteaux et al., 2018a; Ugrankar-Banerjee et al., 2023). Cells with microvilli, such as those that line the lumen of Malpighian tubules (Özyurt et al., 2017), tend to exhibit strong staining for F-actin in the apical (microvilli-rich) region (Kayukawa and Ishikawa, 2009; Meulemans and De Loof, 1990). Similarly, the contractile thin filaments of muscle cells are also rich in F-actin, as seen in muscle surrounding the midgut (Des Marteaux et al., 2018a; Kim et al., 2006). Like F-actin, microtubules can also be found close to the plasma membrane (e.g., fat body in Des Marteaux et al., 2018a), and form internal networks that are important for intracellular transport and cell structure (e.g., neurons in Rolls et al., 2007). Microtubules are also major components of specialized structures such as flagella (e.g., sperm cells in Afzelius et al., 1990). Given the diversity of cytoskeletal arrangements in different tissue types, it is important to understand whether those tissues differentially respond to cold and other environmental influences.

To study the role of the cytoskeleton in freeze tolerance, we worked with an emerging laboratory model organism, *G. veletis*. Fifth instar *G. veletis* become freeze-tolerant when exposed to a 6-week fall-like acclimation of decreasing temperature and photoperiod, and can survive freezing to moderately low temperatures (e.g., −8°C) for several days (Toxopeus et al., 2019c). During acclimation, *G. veletis* differentially express mRNA for actin (downregulation) and tubulin (upregulation) in their fat body tissue (Toxopeus et al., 2019b), along with upregulation of genes encoding proteins that interact with or regulate the cytoskeleton such as supervillin (a member of the villin/gelsolin family; Pestonjamasp et al., 1997), and Jupiter (a tubulin-associated protein; Karpova et al., 2006). Here, we tested the hypothesis that *G. veletis* modify their cytoskeleton in multiple tissues during fall-like acclimation to better protect cytoskeletal structures from the challenges of low temperatures and freezing.

## Materials and Methods

### Insect Rearing and Experimental Design

All experiments were conducted on fifth instar male nymphs of *G. veletis*, reared under summer-like conditions (25°C, 14:10 L:D photoperiod) as previously described (Toxopeus et al., 2019c), or exposed to a 6-week fall-like acclimation that induces freeze tolerance (Adams et al., 2024).

To determine whether acclimation affected the cytoskeleton, we compared tissues from freeze-intolerant crickets haphazardly selected from rearing bins (0 weeks acclimation) and freeze-tolerant crickets that had undergone the full 6 weeks of acclimation (Fig. 1A). We also compared the effect of chilling and freezing on the cytoskeleton in both freeze-intolerant and freeze-tolerant crickets (Fig. 1A). For this second experiment, we compared tissues from control crickets (never chilled or frozen) to crickets were cooled to the same low temperature (−8°C) and either froze or remained unfrozen (“chilled”; Fig. 1B). The chilling treatment was survivable for all crickets, while the freezing treatment was survivable for freeze-tolerant crickets but lethal for freeze-intolerant crickets, allowing us to also examine the effect of lethal and non-lethal freezing. Following all treatments, we stained F-actin and β-tubulin in three tissue types (fat body, Malpighian tubules, and femur muscle) to characterize both qualitative and quantitative changes in the cytoskeleton via methods similar to those used in other insects (Des Marteaux et al., 2018a; Des Marteaux et al., 2018b; Kim and Denlinger, 2009; Kim et al., 2006).

**Figure 1.**
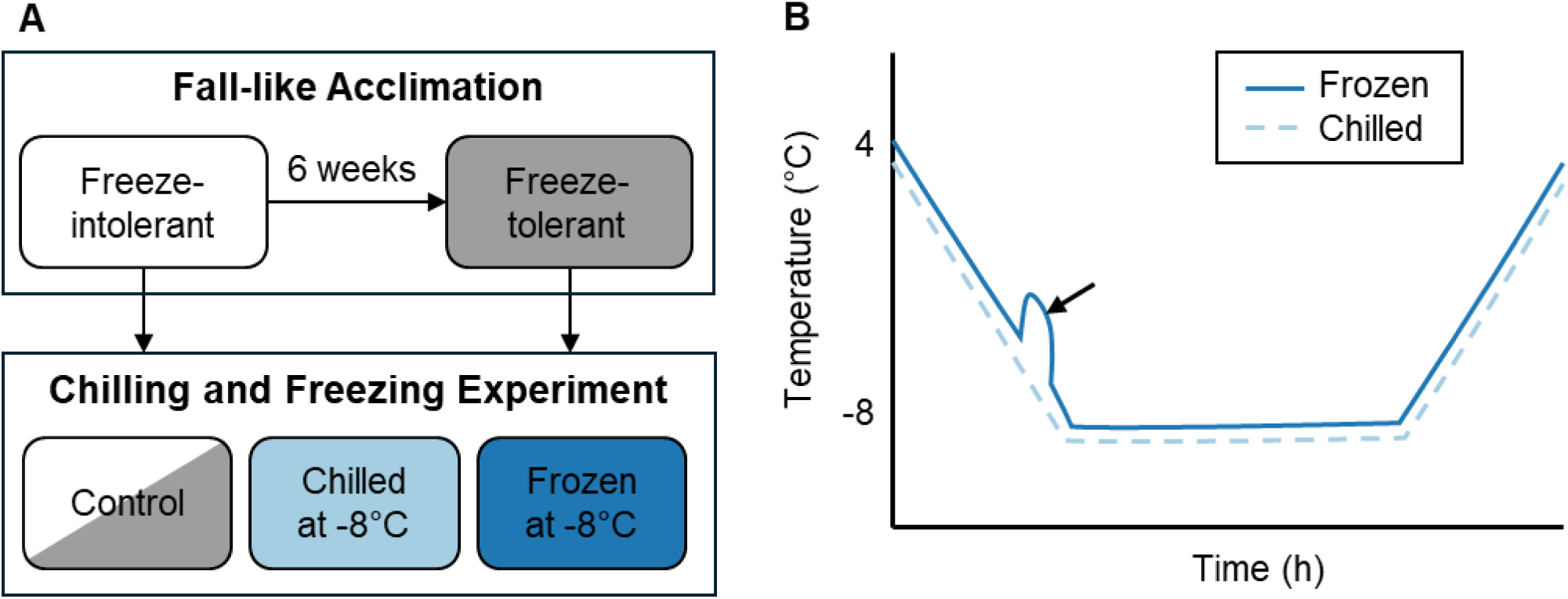
Experimental design to test the effect of acclimation and low temperatures on the cytoskeleton of *Gryllus veletis* tissues. (A) The effect of acclimation was determined by comparing freeze-intolerant (0 weeks acclimation) and freeze-tolerant (6 weeks acclimation) crickets. The effect of chilling and freezing was examined in both freeze-intolerant and freeze-tolerant crickets relative to control (not chilled or frozen) crickets. **(B)** Chilled and frozen crickets were exposed to the same temperatures, but ice nucleation by silver iodide facilitated freezing in half of the crickets, which we detected by the presence of an exotherm (arrow) caused by the process of ice formation.

### Chilling and Freezing Treatments

For freezing and chilling treatments, freeze-tolerant and freeze-intolerant *G. veletis* in 1.7 mL microcentrifuge tubes were placed in a polystyrene foam-insulated aluminum block connected to a ThermoFisher ArcticA25 Immersion Bath Circulator (Adams et al., 2024; Lemay et al., 2024; McIntyre et al., 2023). A drop of silver iodide slurry (1-2 μL) was added to the dorsal side of half of the crickets to facilitate ice nucleation (Sinclair et al., 2015; Toxopeus et al., 2019c). Crickets were exposed to 4°C for 10 min, gradually cooled at −0.25°C min^-1^ to −8°C, held at the target temperature for 90 min, then gradually warmed at 0.25°C min^-1^ to 3°C. To detect freezing (Fig. 1B), the temperature of each cricket was tracked once per second via a T-type constantan-copper thermocouple (Omega Engineering, Norwalk, CT, USA) interfaced with Picolog 6.2.8 software (Pico Technology, Cambridgeshire, UK) via TC-08 USB units.

### Tissue Dissection, Fixation, and Staining

Malpighian tubules, fat body, and femur muscle were dissected in phosphate buffered saline (PBS) following rearing, 6 weeks of acclimation, or within 2 h after a chilling or freezing exposure. Freeze-intolerant crickets collected from 25°C conditions were dissected at room temperature. Freeze-tolerant crickets and any crickets that had been chilled or frozen at sub-zero temperatures were dissected over ice. Tissues were fixed overnight (18-24 h) at 4°C with gentle shaking in 4% paraformaldehyde (PFA) in PBS. Samples were washed at room temperature three times for 10 min each in PBS (if being sectioned or stored prior to staining) or 0.5% Triton X-100 in PBS (PBSTx; if being stained immediately). Samples were stored for up to 21 days at 4°C prior to staining. To improve stain permeation into femur muscle tissue, these samples were embedded in a 3% agarose in PBS block for sectioning into 100 μm slices with a Vibratome® Series 1000 prior to staining.

Fixed tissues underwent immunocytochemistry and simple fluorescent staining in a single protocol to visualize β-tubulin and F-actin, respectively. The following steps were all performed at room temperature. Tissues were permeabilized and blocked for 1 h in 5% normal goat serum (NGS) in PBSTx, incubated for 2 h with 5 μg mL^-1^ mouse anti-β-tubulin primary antibody (E7, Developmental Studies Hybridoma Bank, University of Iowa, Iowa City, IA, USA), and washed three times (10 min) in PBSTx. In each round of staining, one sample per tissue type was used as a negative control with no addition of primary antibody, to account for any possible non-specific staining in the immunocytochemistry process. Tissues were then incubated in darkness overnight (18-24 h) with 0.5% phalloidin conjugated to AlexaFluor 488 (A12379; ThermoFisher Scientific, Toronto, ON, Canada) in PBSTx to stain F-actin, then washed three times (10 min) in PBSTx, and incubated for 4 h with 0.1% goat anti-mouse secondary antibody conjugated to AlexaFluor 647 (A21235; ThermoFisher) in PBSTx. Samples were washed four times (10 min) in PBS prior to arranging whole tissues (Malpighian tubules or fat body) or 100 μm sections (femur muscle) onto glass slides in Fluoromount-G containing DAPI (ThermoFisher) for nuclear staining. Edges of the glass coverslips were sealed with nail polish, and slides were stored in the dark at 4°C for up to 72 h before imaging.

### Imaging and Image Processing

Malpighian tubules, fat body and femur muscle were imaged with an Olympus Fluoview 3000 confocal microscope at 10× (for quantification) or 40× (for high-resolution representation images) objective lens magnification with excitation/emission wavelengths of 405/461 nm for DAPI (nuclei), 488/520 nm for phalloidin (F-actin), and 650/671 nm for the secondary antibody conjugated to AlexaFluor 647 (β-tubulin). Imaging parameters were selected to ensure staining intensity was not saturated in any sample, and there was little to no signal from β-tubulin in negative control samples. Z stacks were taken on the confocal microscope using the entire depth of the tissue in a single field of view. To account for potential effects of tissue thickness on staining intensity, the signal intensity for each target was averaged across all stacks in the Z plane. In FIJI (ImageJ) software (Schindelin et al., 2012), confocal image files were converted to 8-bit, tissue area was traced freehand (to exclude empty background or intruding tissues like tracheoles), and the average pixel intensity for each channel (F-actin [green], β-tubulin [red]) was determined using the measure function (Kim and Denlinger, 2009; Kim et al., 2006). All pixel intensity calculations were standardized to the area of the selected tissue. The staining intensity of β-tubulin was corrected for each sample by subtracting the intensity of the negative control sample.

### Statistical Analysis

All statistical tests were conducted in R version 4.3.3 (R Core Team, 2024). Differences in relative intensity of F-actin and β-tubulin staining between freeze-tolerant and freeze-intolerant crickets were determined by one-tailed Mann-Whitney tests. Differences in relative intensity of F-actin and β-tubulin between control, chilled and frozen crickets were determined by Kruskal-Wallis tests with Dunn-Bonferroni post-hoc tests. Outliers with a 5-fold or greater difference from the mean value of each treatment were removed from statistical analysis.

## Results

### Tissue specific patterns of F-actin and β-tubulin staining

Our staining of F-actin and β-tubulin revealed expected localization of microfilaments and microtubules (respectively) in fat body, Malpighian tubules and femur muscle of *G. veletis* (Fig. 2). In fat body tissue, we found clear staining of microfilaments near the plasma membrane and a network of microtubules within fat body cells (Fig. 2A-C), similar to other insects (Des Marteaux et al., 2018a; Ugrankar-Banerjee et al., 2023; Zheng et al., 2020). In Malpighian tubules, we observed microfilament staining at the apical region of epithelial cells (i.e., the lumen of the Malpighian tubules; Fig. 2D-E), where F-actin is expected to be abundant to support microvilli (Des Marteaux et al., 2018a; Kayukawa and Ishikawa, 2009; Meulemans and De Loof, 1990). We also detected microfilament staining in the thin strands of muscle that spiral around Malpighian tubules (Fig. 2D-E) that are present in some insect species (e.g., cockroaches - Crowder and Shankland, 1972; true bugs - Özyurt et al., 2017), but not others (e.g., flies - Des Marteaux et al., 2018a; Kayukawa and Ishikawa, 2009). We detected faint staining of microtubules in Malpighian tubule epithelial cells (similar to Des Marteaux et al., 2018a), and strong staining in presumptive tracheoles (see Lukinavičius et al., 2018; Özyurt et al., 2017) wrapped around the tubules (Fig. 2D, F). In femur skeletal muscle sections, microfilament staining was localized to contractile filaments (Fig. 2G-H), exhibiting a striated pattern seen in other insect muscle tissue (Des Marteaux et al., 2018a; Kim et al., 2006). Microtubule staining within femur muscle was weak (Fig. 2G, I), similar to midgut muscle of flies (Des Marteaux et al., 2018a), with stronger staining of tubulin in presumptive tracheoles (Fig 2G, I).

**Figure 2.**
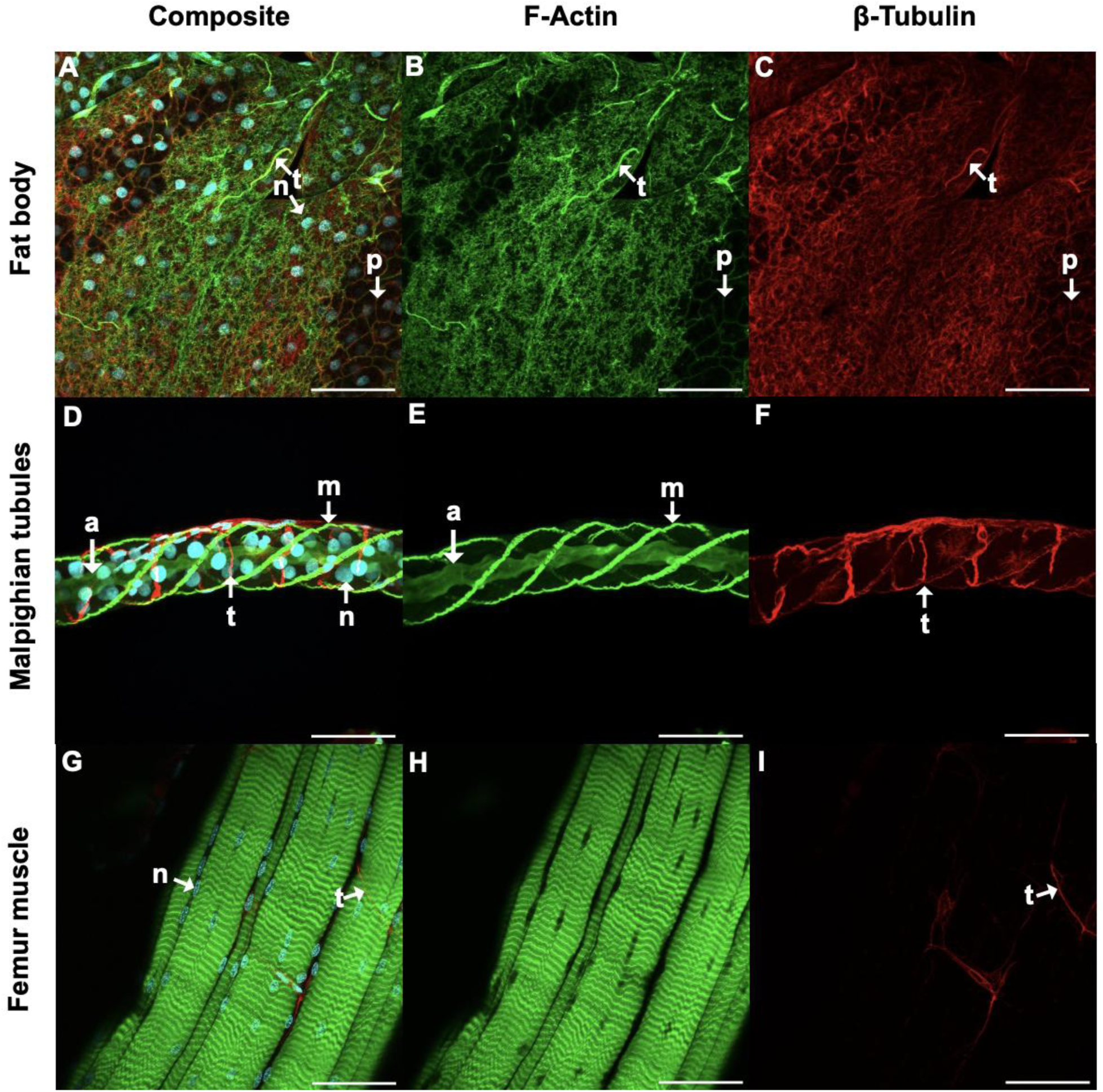
Location of F-actin (green), β-tubulin (red), and nuclei (cyan) staining in *Gryllus veletis* (A-C) fat body, (D-F) Malpighian tubules, and (G-I) femur muscle. Staining details are in the methods. Images were obtained via confocal microscopy and maximum-intensity projections of the Z-stacks are shown here to optimize viewing of stain location. n, nucleus; t, tracheole; p, plasma membrane; m, muscle; a, apical region of epithelial cells. Scale bar is 50 μm in all panels.

### F-actin, but not β-tubulin, staining intensity is increased in tissues of freeze-tolerant crickets

While F-actin staining intensity was affected by the acclimation that induces freeze tolerance in *G. veletis*, β-tubulin staining intensity was not. There was an increase in relative intensity of F-actin staining in fat body (Fig. 3A-C; *W* = 124, *P* = 0.008) and Malpighian tubules (Fig. 3D-F; *W* = 67, *P* < 0.001) in freeze-tolerant crickets compared to freeze-intolerant crickets, suggesting that acclimation caused an increase in F-actin abundance in these tissues. In Malpighian tubules, the increase in F-actin intensity was generally associated with the apical portion of epithelial cells, rather than the muscle fibres around the tubule exterior (Fig. E-F). F-actin staining in femur muscle remained unchanged during acclimation (Fig 3G-I; *W* = 111, *P* = 0.385). In addition, β-tubulin intensity did not change with acclimation in *G. veletis* fat body (Fig. 4A-C, *W* = 207, *P* = 0.717), Malpighian tubules (Fig. 4D-F; *W* = 34, *P* = 0.196), or femur muscle (Fig. 4G-I; *W* = 20, *P* = 0.126). There were no clear qualitative changes in staining (e.g., localization) for either F-actin (Fig. 3) or β-tubulin (Fig. 4) for any tissue during acclimation.

**Figure 3.**
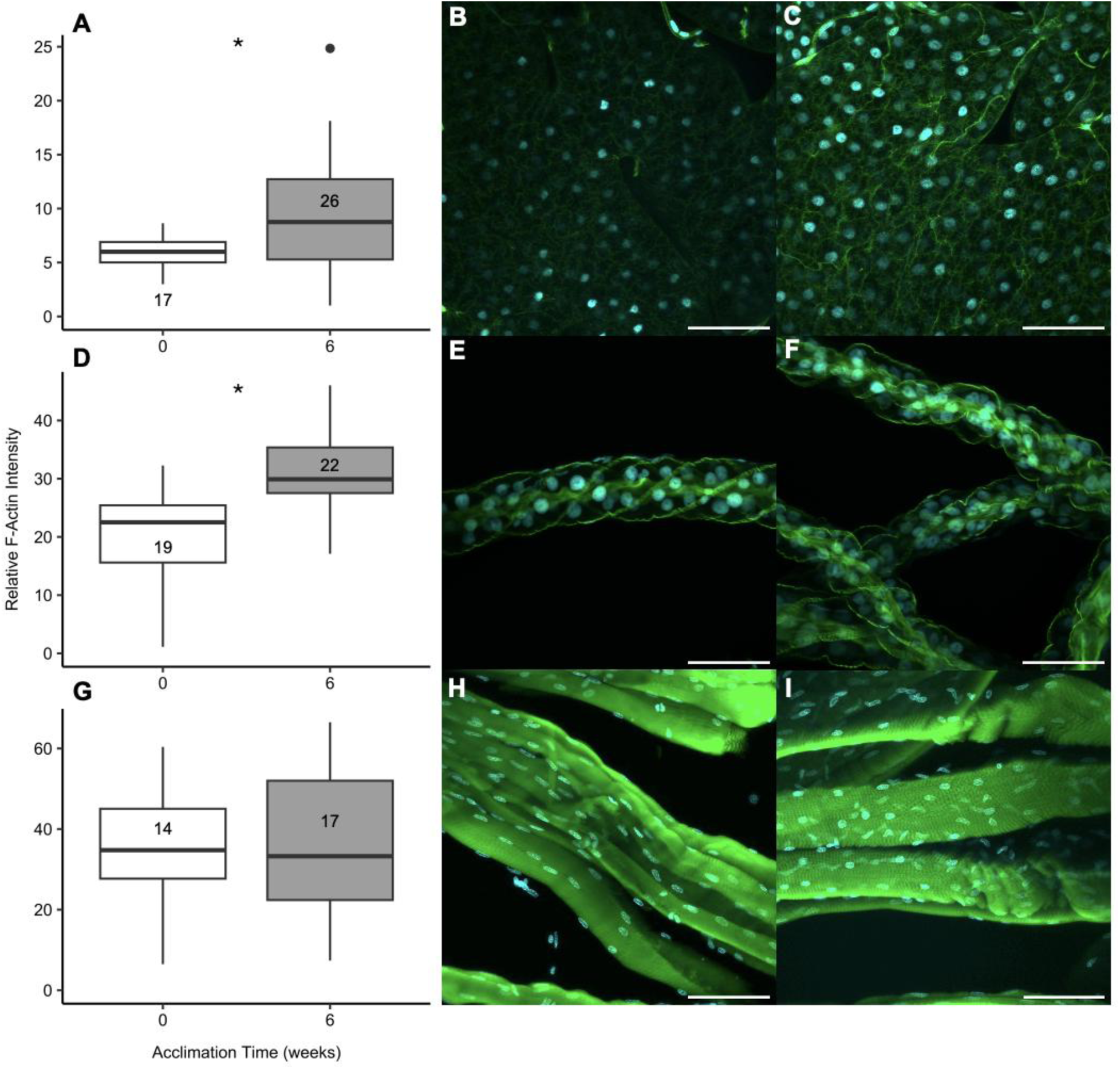
F-actin staining intensity and location in (A-C) fat body, (D-F) Malpighian tubules, and (G-I) femur muscle from *Gryllus veletis* before (0 weeks) and after (6 weeks) the fall-like acclimation that induces freeze-tolerance. F-actin (green) and nuclei (cyan) were stained as described in the Methods. **(A, D, G)** Numbers within or below each boxplot depict sample size (number of crickets) for intensity quantification. The upper quartile (box top), median (thick centre line), lower quartile (box bottom), and the minimum and maximum values within 1.5 times the inter-quartile range (whiskers) are shown. Black dots represent outliers. Asterisks indicates statistical increase (*P* < 0.05) in F-actin staining intensity between freeze-tolerant and freeze-intolerant crickets, as determined by Mann-Whitney tests. Images with representative F-actin staining intensity (close to the mean) are shown for **(B, E, H)** 0-week (freeze-intolerant) and **(C, F, I)** 6-week (freeze-tolerant) acclimated crickets. Scale bar is 50 μm in all panels.

**Figure 4.**
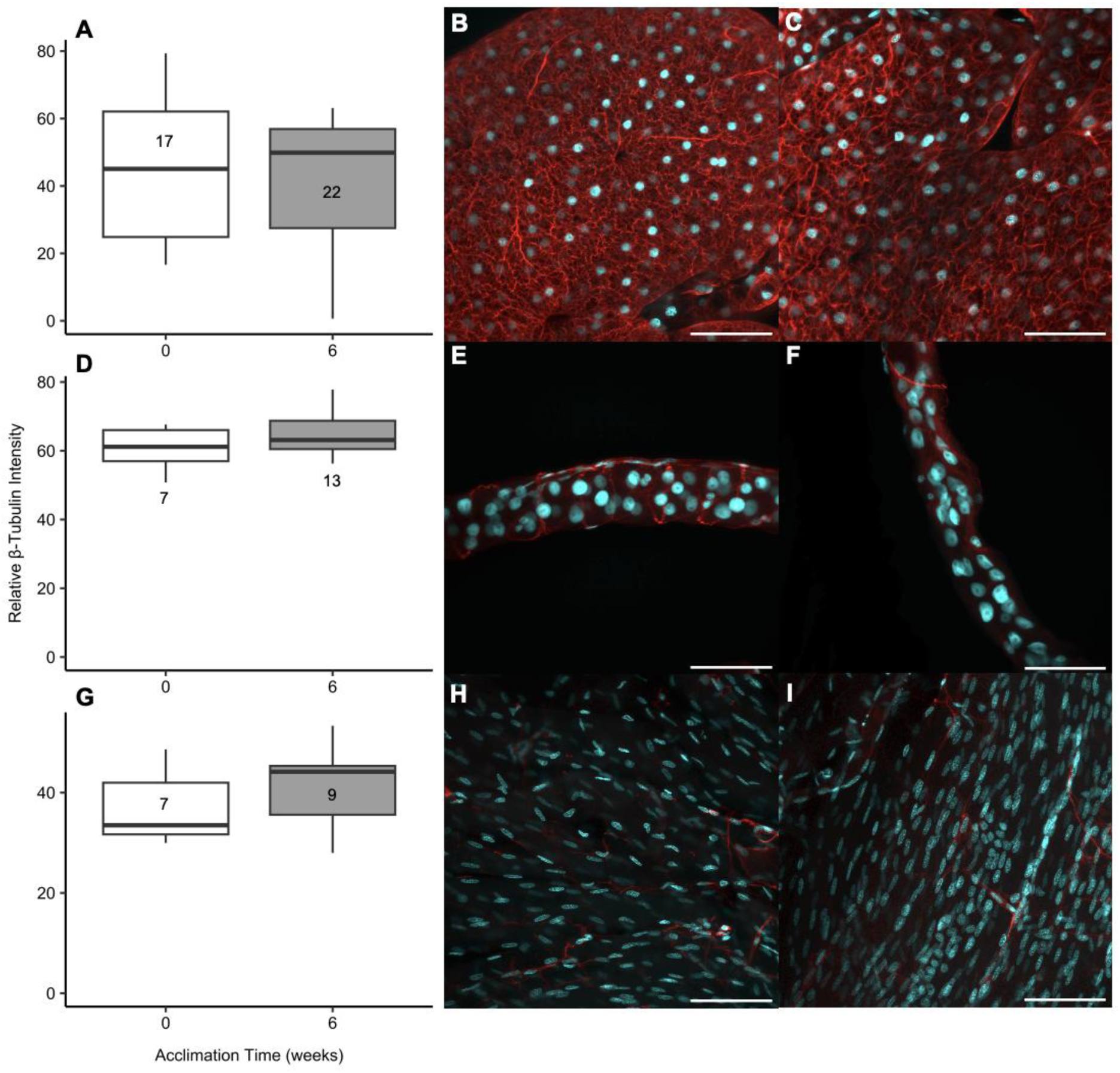
β-tubulin staining intensity and location in (A-C) fat body, (D-F) Malpighian tubules, and (G-I) femur muscle from *Gryllus veletis* before (0 weeks) and after (6 weeks) the fall-like acclimation that induces freeze-tolerance. β-tubulin (red) and nuclei (cyan) were stained as described in Methods. (A, D, G) Numbers within or below each boxplot depict sample size (number of crickets) for intensity quantification. The upper quartile (box top), median (thick centre line), lower quartile (box bottom), and the minimum and maximum values within 1.5 times the inter-quartile range (whiskers) are shown. There were no statistical differences (*P* > 0.05) in β-tubulin staining intensity between freeze-tolerant and freeze-intolerant crickets, as determined by Mann-Whitney tests. Images with representative β-tubulin staining intensity (close to the mean) are shown for **(B, E, H)** 0-week (freeze-intolerant) and **(C, F, I)** 6-week (freeze-tolerant) acclimated crickets. Scale bar is 50 μm in all panels.

### Freezing causes tissue-specific changes in F-actin and β-tubulin staining intensity

F-actin staining was affected by freezing in the fat body of freeze-intolerant crickets, freeze-tolerant cricket F-actin and β-tubulin were affected by chilling or freezing in Malpighian tubules, and neither cytoskeletal element was impacted by our treatments in femur muscle. In fat body (Fig. 5A-C), F-actin staining intensity decreased in freeze-intolerant crickets following freezing but not chilling (*H* = 6.000, *P* = 0.049), while no changes were observed in freeze-tolerant crickets (*H* = 2.052, *P* = 0.358). In the Malpighian tubules (Fig. 5D), F-actin intensity was unaffected by freezing or chilling in freeze-intolerant crickets (*H* = 1.502, *P* = 0.472), but decreased in the Malpighian tubules of freeze-tolerant crickets that had been chilled (*H* = 8.582, *P* = 0.014; Fig. 5D-F). In femur muscle (Fig. 5G-I), F-actin of both freeze-intolerant crickets (*H* = 4.068, *P* = 0.131) and freeze-tolerant crickets (*H* = 0.729, *P* = 0.694) were unaffected by chilling or freezing. Similar to the acclimation results, β-tubulin staining intensity generally remained stable following chilling and freezing in fat body (Fig. 6A-C; freeze-intolerant: *H* = 0.130, *P* =0.937; freeze-tolerant: *H* = 1.125, *P*=0.570), Malpighian tubules (Fig. 6D-F; freeze-intolerant: *H* = 0.795, *P* = 0.672), and femur muscle (Fig. 6G-I; freeze-intolerant: *H* = 0.612, *P* = 0.736; freeze-tolerant: *H* = 1.453, *P* = 0.483). However, freezing was associated with decreased β-tubulin staining intensity in freeze-tolerant crickets (Fig. 6D; *H* = 11.018, *P* = 0.004). There were no clear qualitative changes in staining (e.g., localization) associated with chilling or freezing for either F-actin (Fig. 5) or β-tubulin (Fig. 6) for any tissue.

**Figure 5.**
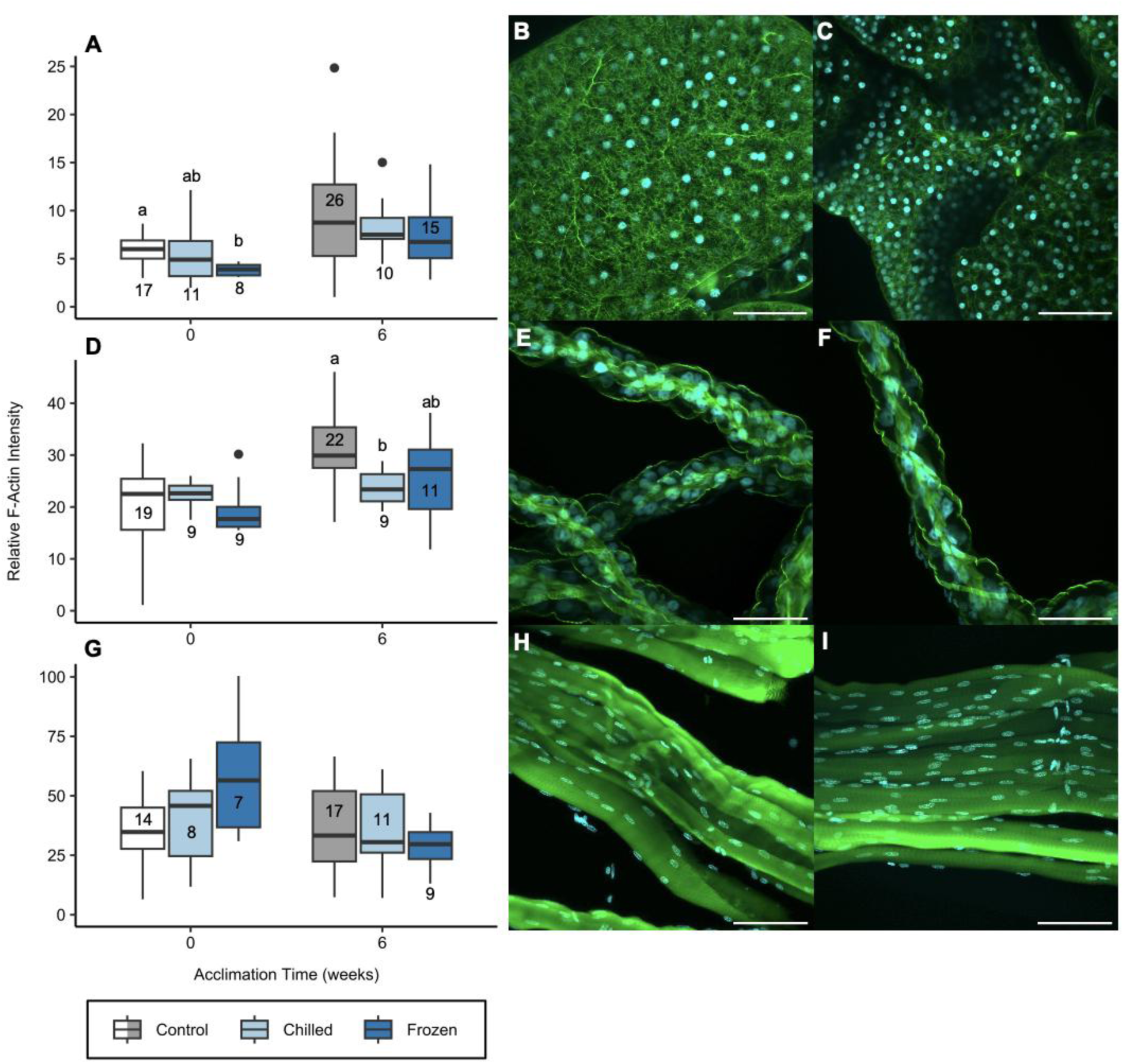
F-actin staining intensity and location in (A-C) fat body, (D-F) Malpighian tubules, and (G-I) femur muscle from unstressed (control; white or gray), chilled (light blue), or frozen (dark blue) *Gryllus veletis* that were freeze-intolerant (0 weeks acclimation) or freeze-tolerant (6 weeks acclimation). F-actin (green) and nuclei (cyan) were stained as described in the Methods. **(A, D, G)** Numbers within or below each boxplot depict sample size (number of crickets) for intensity quantification. The upper quartile (box top), median (thick centre line), lower quartile (box bottom), and the minimum and maximum values within 1.5 times the inter-quartile range (whiskers) are shown. Black dots represent outliers. Different letters indicate statistical differences (*P* < 0.05) in F-actin staining intensity between treatments within the freeze-intolerant or freeze-tolerant crickets, as determined by Kruskal-Wallis with Dunn-Bonferroni post-hoc tests. Images with representative F-actin staining intensity (close to the mean) are shown for treatments of interest (e.g., statistically different), including fat body from freeze-intolerant **(B)** control and **(C)** frozen crickets, Malpighian tubules from freeze-tolerant **(E)** control and **(F)** chilled crickets, and femur muscle from freeze-intolerant **(H)** control and **(I)** frozen crickets. Scale bar is 50 μm in all panels.

**Figure 6.**
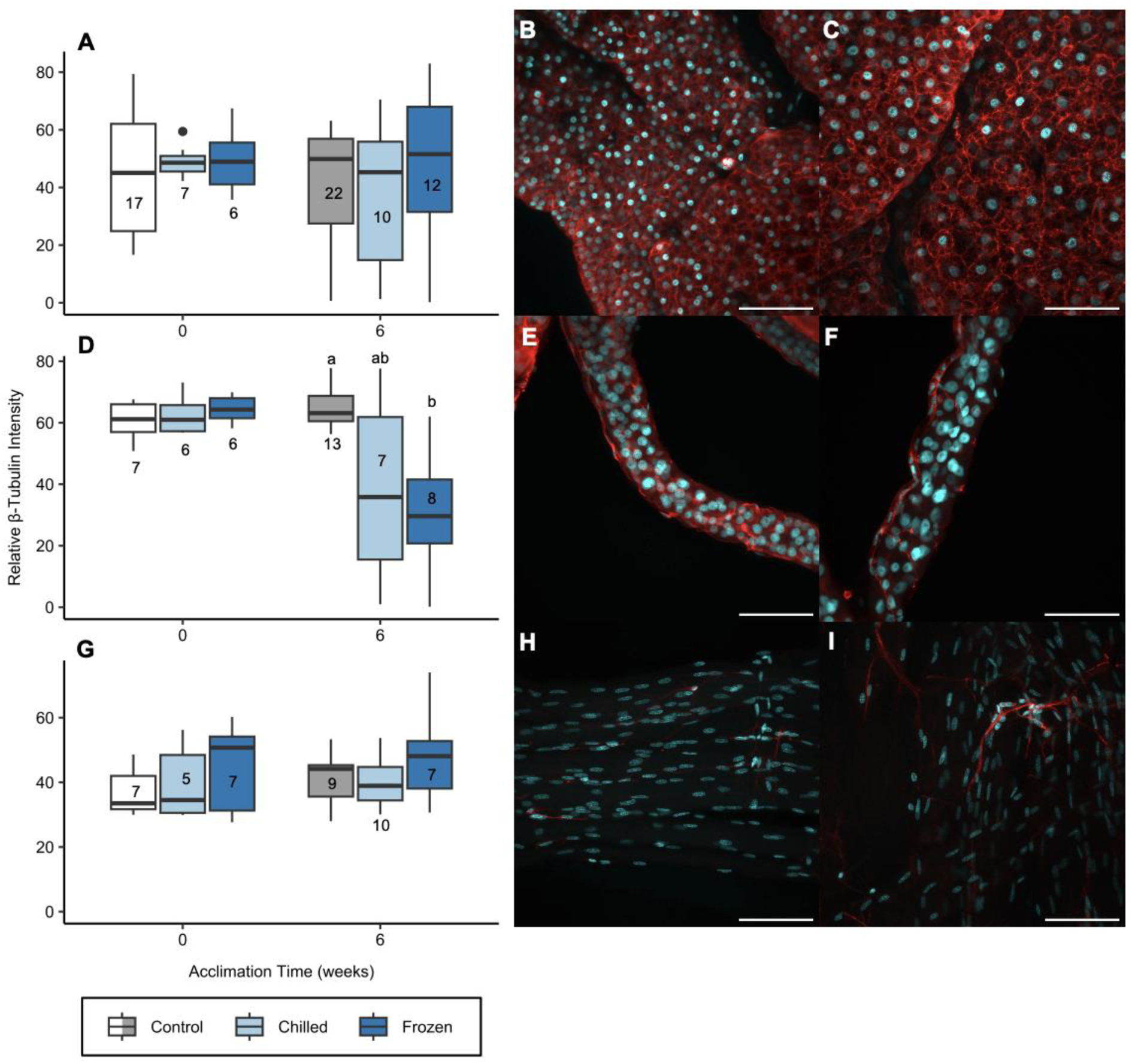
β-tubulin staining intensity and location in (A-C) fat body, (D-F) Malpighian tubules, and (G-I) femur muscle from unstressed (control; white or gray), chilled (light blue), or frozen (dark blue) *Gryllus veletis* that were freeze-intolerant (0 weeks acclimation) or freeze-tolerant (6 weeks acclimation). β-tubulin (red) and nuclei (cyan) were stained as described in Methods. **(A, D, G)** Numbers within or below each boxplot depict sample size (number of crickets) for intensity quantification. The upper quartile (box top), median (thick centre line), lower quartile (box bottom), and the minimum and maximum values within 1.5 times the inter-quartile range (whiskers) are shown. Black dots represent outliers. Different letters indicate statistical differences (*P* < 0.05) in β-tubulin staining intensity between treatments within the freeze-intolerant or freeze-tolerant crickets, as determined by Kruskal-Wallis with Dunn-Bonferroni post-hoc tests. Images with representative β-tubulin staining intensity (close to the mean) are shown for frozen crickets from both the **(B, E, H)** freeze-intolerant and **(C, F, I)** freeze-tolerant groups. Scale bar is 50 μm in all panels.

## Discussion

Our study is one of few that has examined the cytoskeleton in freeze-tolerant insects, and the first to do so in *G. veletis*. We saw tissue-specific staining of both microfilaments and microtubules that aligned well with what is known about cytoskeletal structure of fat body, Malpighian tubules, and muscle in other insects (see Results). As hypothesized, there was a clear increase in microfilament abundance in fat body and Malpighian tubules (but not femur muscle) during the acclimation that induces freeze tolerance. Conversely, we did not see a change in abundance of β-tubulin, nor substantial changes in the arrangements of microtubules during acclimation. Consistent with our hypothesis, freeze-tolerant crickets maintained their cytoskeletal structure in fat body following chilling and freezing to subzero temperatures (c. −8°C), while freeze-intolerant crickets exhibited microfilament damage in fat body tissue after freezing but not chilling. Contrary to our hypothesis, the microfilament and microtubule cytoskeleton of Malpighian tubules in freeze-tolerant crickets seemed damaged by chilling or freezing. Our results suggest that cytoskeletal modification may be an important freeze tolerance adaptation to preserve cell structure in fat body, but not all tissues.

### Acclimation alters the actin cytoskeleton of freeze-tolerant crickets

The tissue-specific increase in microfilament (F-actin) abundance associated with freeze tolerance in *G. veletis* may be important for supporting plasma membrane structure. Others have shown a close relationship between the actin cytoskeleton and plasma membrane shape/projections (Des Marteaux et al., 2018a; Ragoonanan et al., 2010; Ugrankar-Banerjee et al., 2023). Cold acclimation also stimulates F-actin accumulation in rectal epithelial cells of *G. pennsylvanicus* (Des Marteaux et al., 2018b), suggesting phenotypic plasticity of the cytoskeleton is important in preparing for cold stress. An increase in microfilament abundance during acclimation of *G. veletis* may therefore help support the plasma membranes of fat body cells in advance of stressful low temperature exposure. However, the increase in microfilament abundance during acclimation in Malpighian tubules did not facilitate maintenance of the actin cytoskeleton microfilament abundance after chilling or freezing, which we discuss below. We speculate that microfilament arrangement and abundance did not change in *G. veletis* femur muscle because these microfilaments are part of sarcomere contractile units and stabilized by many other proteins (Pollard, 2016).

We hypothesize that the mechanism of increased microfilament abundance in *G. veletis* fat body and Malpighian tubules is likely driven by a rearrangement of an existing pool of G-actin into F-actin, rather than the synthesis of new actin proteins. Because phalloidin stains only F-actin (not the monomeric G-actin), we have no measurements of G-actin abundance in this study.

However, previous transcriptomic work in *G. veletis* has shown that all actin isoforms are either not differentially expressed or are downregulated in fat body tissue during fall-like acclimation (Toxopeus et al., 2019b), suggesting that synthesis of new actin protein is unlikely. Others have also shown that cytoskeletal filaments (polymers) can increase or decrease while the abundance of cytoskeletal monomers remains stable (Li and Moore, 2020; Pokorná et al., 2004). Similar to other insect species (Des Marteaux et al., 2017; Kayukawa and Ishikawa, 2009), cytoskeletal regulators (e.g., Supervillin, Jupiter) are upregulated during acclimation in *G. veletis* (Toxopeus et al., 2019b), which may facilitate organization of more microfilaments. To determine the mechanisms underlying microfilament accumulation during acclimation, additional work such as semi-quantitative Western blotting (to measure actin monomer abundance; e.g., Adams et al., 2024; King et al., 2013) and manipulation of actin cytoskeleton regulators such as Supervillin (e.g., via RNAi; Toxopeus, 2018) could be conducted.

Our β-tubulin staining results suggest that freeze tolerance is not associated with microtubule accumulation in *G. veletis*. While our staining method did not distinguish between β-tubulin monomers (single proteins) and β-tubulin incorporated into microtubules, we observed clear filamentous structures (microtubules) in fat body that seemed unaffected by our fall-like acclimation. Our staining results are unexpected, given the previously-described upregulation of tubulin mRNA in fat body during fall-like acclimation *G. veletis* (Toxopeus et al., 2019b).

However, our results are similar to studies in other insects, including freeze-tolerant *C. costata* that undergo very few modifications to their microtubule arrangement in multiple tissues during acclimation (Des Marteaux et al., 2018a), and *Culex pipiens* mosquitoes that exhibit decreased microtubule abundance in their flight muscle in association with dormancy (diapause) and increased cold tolerance (Kim and Denlinger, 2009). Due to the subtle staining of tubulin in *G. veletis* Malpighian tubules and femur muscle, it may be worth investigating microtubule structure in these tissues via other methods (e.g., Li and Moore, 2020; Lukinavičius et al., 2018; Wang et al., 2019) to expand our understanding of how cytoskeletal structure can change with acclimation.

### Ice, not low temperatures, causes damage to the actin cytoskeleton of freeze-intolerant crickets

Chilling did not appear to affect microfilament and microtubule abundance and arrangement in freeze-intolerant crickets, suggesting that either subzero temperatures do not depolymerize the cytoskeleton in these crickets, or that they can successfully repolymerize cytoskeletal filaments during rewarming. While some studies have shown depolymerization of microfilaments or microtubules during or after exposure to low temperatures (Des Marteaux et al., 2018b; Kim and Denlinger, 2009; Pokorná et al., 2004), others have shown that microfilaments may resist depolymerization in the cold (Li and Moore, 2020; Müllers et al., 2019). Our chilling treatment included slow ramping (0.25°C min^-1^) during cooling to −8°C and warming to 4°C, which may have decreased the intensity of the cold shock (minimizing depolymerization) and provided some time for repolymerization during warming (compare to Bartolo and Carter, 1991; Li and Moore, 2020; Pokorná et al., 2004). More intense cold treatments with lower temperatures, longer durations, or sharper temperatures transitions (e.g., Kayukawa and Ishikawa, 2009; Kim and Denlinger, 2009) may cause more cytoskeletal damage during or after chilling than we observed in our study. However, the minimal impact of chilling on freeze-intolerant (unacclimated) *G. veletis* in the present study suggests that the dominant effect of freezing on the cytoskeleton of fat body in these crickets was caused by ice, rather than low temperature itself.

In freeze-intolerant crickets that were frozen and thawed, the abundance and arrangement of microfilaments but not microtubules changed, suggesting that lethal freezing has an impact on the actin cytoskeleton. In particular, the fat body of frozen freeze-intolerant crickets showed a decrease in F-actin abundance, showing low resilience to the effects of ice formation. Plasma membrane integrity is compromised in fat body from frozen freeze-intolerant *G. veletis* (Toxopeus et al., 2019c), which could allow cytoskeletal subunits (e.g., G-actin) to leak out of the cell (e.g., Ragoonanan et al., 2010). In addition, it is hypothesized that ice formation occurs extracellularly (in the gut or hemolymph) in freeze-tolerant *G. veletis* (Toxopeus and Sinclair, 2018; Toxopeus et al., 2019c), but intracellular ice formation may occur in the fat body of freeze-intolerant crickets, which could cause physical damage to actin microfilaments (e.g., Müllers et al., 2019). While we did not see clear damage to the cytoskeleton of freeze-intolerant crickets in Malpighian tubules or femur muscle, several other studies have shown that permanent cytoskeletal damage occurs several hours after a cold or freezing stress (Bartolo and Carter, 1991; Kayukawa and Ishikawa, 2009; Müllers et al., 2019). To fully characterize the cytoskeletal damage caused by lethal freezing in multiple tissues, additional work that examines both microtubules and microfilaments during a recovery period post-thaw would be required.

### The actin cytoskeleton is robust to freezing in select tissues of freeze-tolerant crickets

Congruent with our predictions, the cytoskeleton of freeze-tolerant (acclimated) cricket fat body was robust to the effects of freezing, suggesting that the physiological and cellular changes during acclimation help preserve cytoskeletal structure in this tissue during a non-lethal freeze treatment. Our results are similar to others who have shown that acclimation promotes increased cytoskeletal stability during or after cold and freezing stress (Bartolo and Carter, 1991; Des Marteaux et al., 2018b; Kayukawa and Ishikawa, 2009). In addition, other work in freeze-tolerant insects has shown surprisingly little impact of freezing on cytoskeletal structure (Des Marteaux et al., 2018a). Additional work that conducts live cell imaging during cooling and freezing cricket tissues *ex vivo* (e.g., Li and Moore, 2020; Müllers et al., 2019) would increase our understanding of whether any cytoskeletal changes occur in tissues that appear to have a stable cytoskeleton after low temperature exposures.

However, in Malpighian tubules of freeze-tolerant crickets both chilling and freezing negatively affected microfilaments and microtubules (respectively), despite having no impact on this tissue in freeze-intolerant crickets. We speculate that acclimation caused changes in Malpighian tubules that made them more vulnerable to chilling or freezing stress. Malpighian tubules are an important ionoregulatory tissue (Des Marteaux et al., 2017; Nocelli et al., 2016) that may be expending additional energy during acclimation to maintain ion and water balance and avoid chill coma (Overgaard and MacMillan, 2017), especially at temperatures near 0°C at the end of acclimation (Adams et al., 2024). This may decrease the amount of energy available for maintaining or repairing cytoskeletal structure following freezing or chilling stress. Future work could test whether Malpighian tubules require more time to reestablish cytoskeletal structures after nonlethal freezing or chilling events than other tissues of freeze-tolerant crickets.

## Conclusions

Our work in the naturally freeze-tolerant *G. veletis* expands our understanding of how the cytoskeleton may support cell survival during and following ice formation. In particular, we showed that F-actin accumulation in fat body of freeze-tolerant crickets appears to support maintenance of microfilament structure during a moderate and survivable freeze treatment. Conversely, changes to Malpighian tubules during acclimation were correlated with reduced cytoskeletal stability during freezing and chilling. Interestingly, in freeze-intolerant crickets, chilling at subzero temperatures had no short-term impact on F-actin, suggesting that the freeze-induced damage to microfilaments was caused by ice rather than low temperatures themselves. We did not observe any notable trends in microtubule biology of fat body, Malpighian tubules, or femur muscle during acclimation. Interesting questions for future research include what happens to the cytoskeleton during (rather than simply after) chilling and freezing, whether any cytoskeletal changes occur during post-thaw recovery periods, and the mechanisms (e.g., cytoskeletal regulators) that support cytoskeletal restructuring in freeze-tolerant insects.

## Acknowledgements

The authors would like to thank V.E. Adams, A.L. Gough, and the StFX Animal Care Facility for help rearing and processing crickets, and to R. Cozzi, W.S. Marshall, L. McIntyre, and R.C. Wyeth for help or advice with immunocytochemistry experiments, including confocal imaging and analysis.

## Data and Resource Availability

All data and code used to analyze data are available in the following Zenodo repository (van Oirschot and Toxopeus, 2024): https://doi.org/10.5281/zenodo.14152035

